# Chemical constituents involved in e-cigarette, or vaping product use-associated lung injury (EVALI)

**DOI:** 10.1101/2020.01.14.905539

**Authors:** Thivanka Muthumalage, Michelle R. Friedman, Matthew D. McGraw, Alan E. Friedman, Irfan Rahman

**Author notes:** Correspondence should be addressed to: Irfan Rahman, Ph.D., Department of Environmental Medicine, University of Rochester Medical Center, Box 850, 601 Elmwood Avenue, Rochester 14642, NY, USA. The authors declare they have no actual or potential competing financial interests.

## Abstract

**Background:** The Centers for Disease Control (CDC) declared e-cigarette (e-cig), or vaping product use-associated lung injury (EVALI) a national outbreak due to the high incidence of emergency department admissions and deaths. Investigators have identified vitamin E acetate (VEA) as the plausible cause for EVALI, based on compounds found in bronchoalveolar lavage fluid.

**Objectives:** We defined the chemical constituents present in e-cig cartridges associated with EVALI and compared constituents to medical-grade and cannabidiol (CBD) containing cartridges.

**Methods:** We measured chemicals and elemental metals in e-liquid and vapor phases of e-cig counterfeit cartridges by Gas Chromatography (GC) and Mass Spectrometry (MS), EPA method TO-15 by GCMS, and ICP-MS analysis.

**Results:** We have identified chemical constituents in e-cig vaping tetrahydrocannabinol (THC)-containing counterfeit cartridges compared to medical-grade and cannabidiol (CBD) containing cartridges. Apart from VEA and THC, other potential toxicants correlated with EVALI included solvent-derived hydrocarbons, silicon conjugated compounds, various terpenes, pesticides/plasticizers/polycaprolactones, and metals. These chemicals are known to cause symptoms, such as cough, shortness of breath or chest pain, nausea, vomiting or diarrhea, fatigue, fever, or weight loss, all symptoms presenting in patients with EVALI.

**Conclusion:** This study provides insights into understanding the chemical-induced disease mechanism of acute lung injury.

## Introduction

Electronic cigarette (e-cig) use or vaping gained popularity in the past decade, especially among youth, due to the availability of many enticing flavors and devices. In August 2019, CDC started reporting e-cigarette, or vaping, product use-associated lung injury (EVALI) cases. The majority of patients with EVALI report vaping tetrahydrocannabinol (THC)-containing counterfeit street e-cig products. By December 2019, there have been over 2,500 hospitalized EVALI patients, most of whom are men younger than 35 years of age (Hartnett et al. 2019). Patients hospitalized with EVALI have manifested several radiological imaging patterns, such as lipoid pneumonia, eosinophilic pneumonia, and chemical damage to the lung tissue (Henry et al. 2019; Lu et al. 2020). Patients diagnosed with this illness have reported symptoms, such as cough, shortness of breath, or chest pain, nausea, vomiting, or diarrhea fatigue, fever, or weight loss (Kalininskiy et al 2019).

Counterfeit street cartridges are produced by using cutting agents, such as medium-chain triglyceride (MCT) oil, vitamin E (tocopherol) acetate (VEA), and seized drugs, such as butane hash oil (Layden et al. 2019). Currently, vitamin E acetate (VEA) has been identified as a key agent involved in EVALI (Blount et al. 2019b). However, VEA would not impose chemical-induced pneumonia, as VEA is protective in the lung upon aerosolization (Hybertson et al. ; Wang et al.). We hypothesized that, while VEA may be present in cartridges and BALF of patients with EVALI, there may be other constituents in these e-cig THC-containing cartridges that can act singularly, or in concert with other constituents, to induce augmented synergistic injuries to the lung.

Thus, in this study, we performed a screening for chemical constituents in liquid and vapor phases from counterfeit street cartridges. We also identified and compared the chemical constituents in CBD-containing cartridges and medical-grade vaping cartridges. Determining the potential culprits in each category may help to understand the underlying mechanism of lung injury associated with e-cigarette or vaping use.

## Methods

### Ethics statement on biosafety approval

All experiments performed in this study were approved and in accordance with the University of Rochester Institutional Biosafety Committee.

### Counterfeit street/patient, CBD-containing, and medical-grade cartridges

E-cig vaping counterfeit (bootleg/street), CBD-containing, and/or medical-grade cartridges (Columbia Care, TheraCeed, New York) were obtained from local vape shops and users/patients. The counterfeit street cartridge brands recovered from patients with EVALI (Kalininskiy et al 2019) included Dank Vape, ROVE, Super G Cookies, Runtz, Chronic Billy Kimber, Exotic (Paris OG), Jungle boys, Most Coast, Chronic Gushers, Chronic Gruntz, Jungle skittles, TKO, Clear Chronic, Exotic Carts, and Smart Carts. No manufacturers’ details were given on cartridges or packing materials. The tested variants of cannabis oil included *sativa, indica,* and *hybrid*. Medical-grade cartridge was used as a known positive comparison control.

### Screening for chemical constituents in e-liquids by GCMS

Chemical constituents in counterfeit or patient-provided (n=38), CBD-containing (n=4), and medical-grade (positive) e-cig vaping cartridges were determined by Gas Chromatography (GC) and Mass Spectrometry (MS).

E-cigarette liquids were diluted 100X into spectral grade methanol and injected into the GCMS detector (Agilent 7890A gas chromatograph with 5975 MSD detector). The system used helium as the carrier gas, flowing 1.2mL/min through an Agilent Technologies column (HP-5MS, 30m x 0.250mm, 0.25 μm, 19091S-433). The oven program initiated at 45°C for 1 minute, ramped to 285°C over 7 minutes at 10°C/min, ramped to 300°C for 10 minutes at 10°C/min, and finally ramped a third time to 325°C over 5 minutes at 5°C/min. The total run time was 53.5 minutes, and the injection volume was 1μL from a 10μL syringe. The samples were analyzed by electron impact ionization in positive ion mode with a mass range of 50-550m/z, with the source temperature at 230°C and the quadrupole at 150°C. Data analysis was performed using Agilent ChemStation software, with ion scans searched against the NIST database for identification.

Chemicals commonly found in each category (counterfeit, CBD-containing, and medical-grade) of e-cig cartridges, but not found in other categories were identified and classified into terpenes, silicon compounds, pesticide constituents, flavor additives, cannabinoids, plasticizers/ polycaprolactones (PCP)/drugs/manufacturing agents, humectants/oil/plant components, vitamin and conjugates, and miscellaneous constituents **(Table 1)**.

**Table 1:**
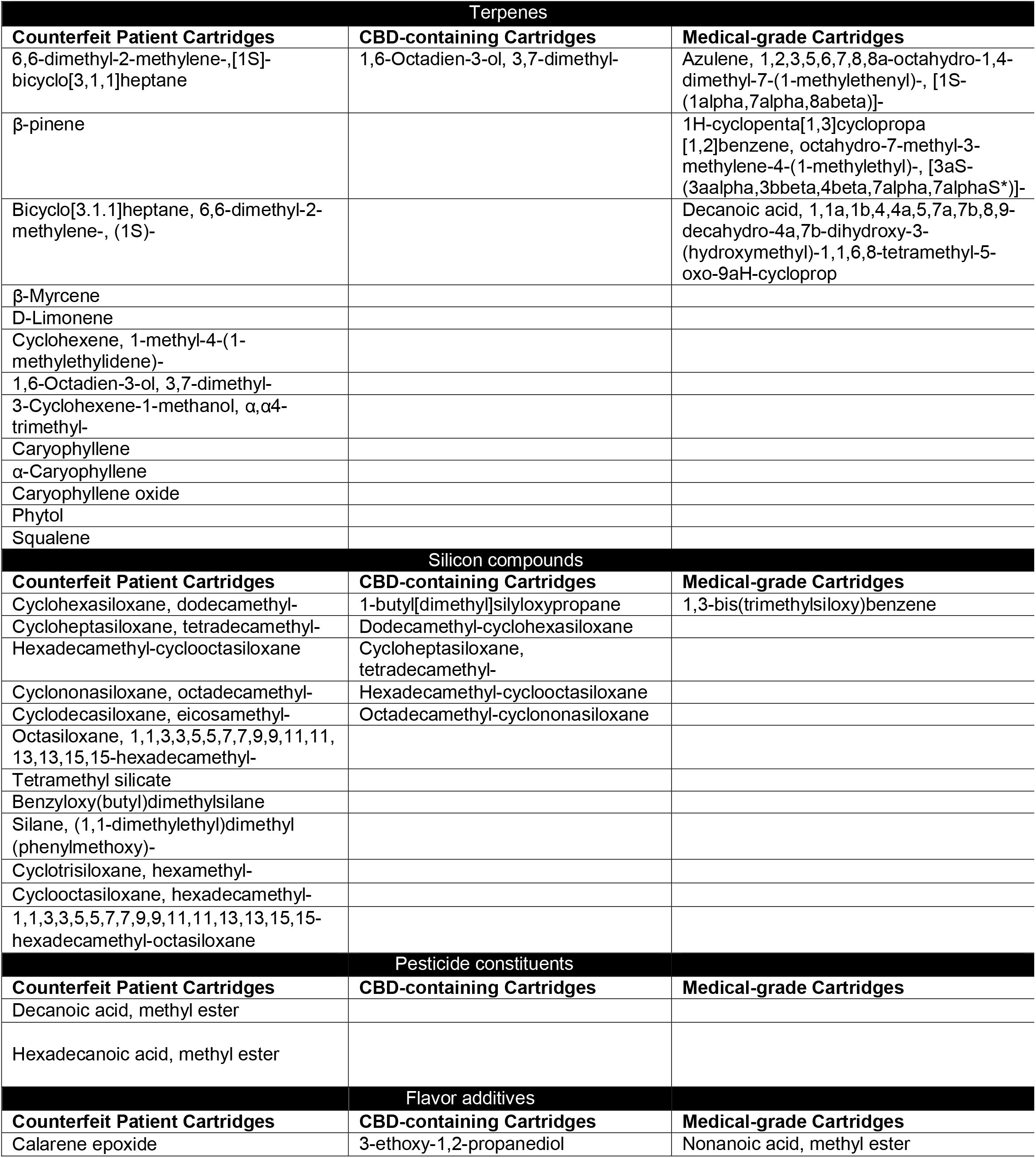
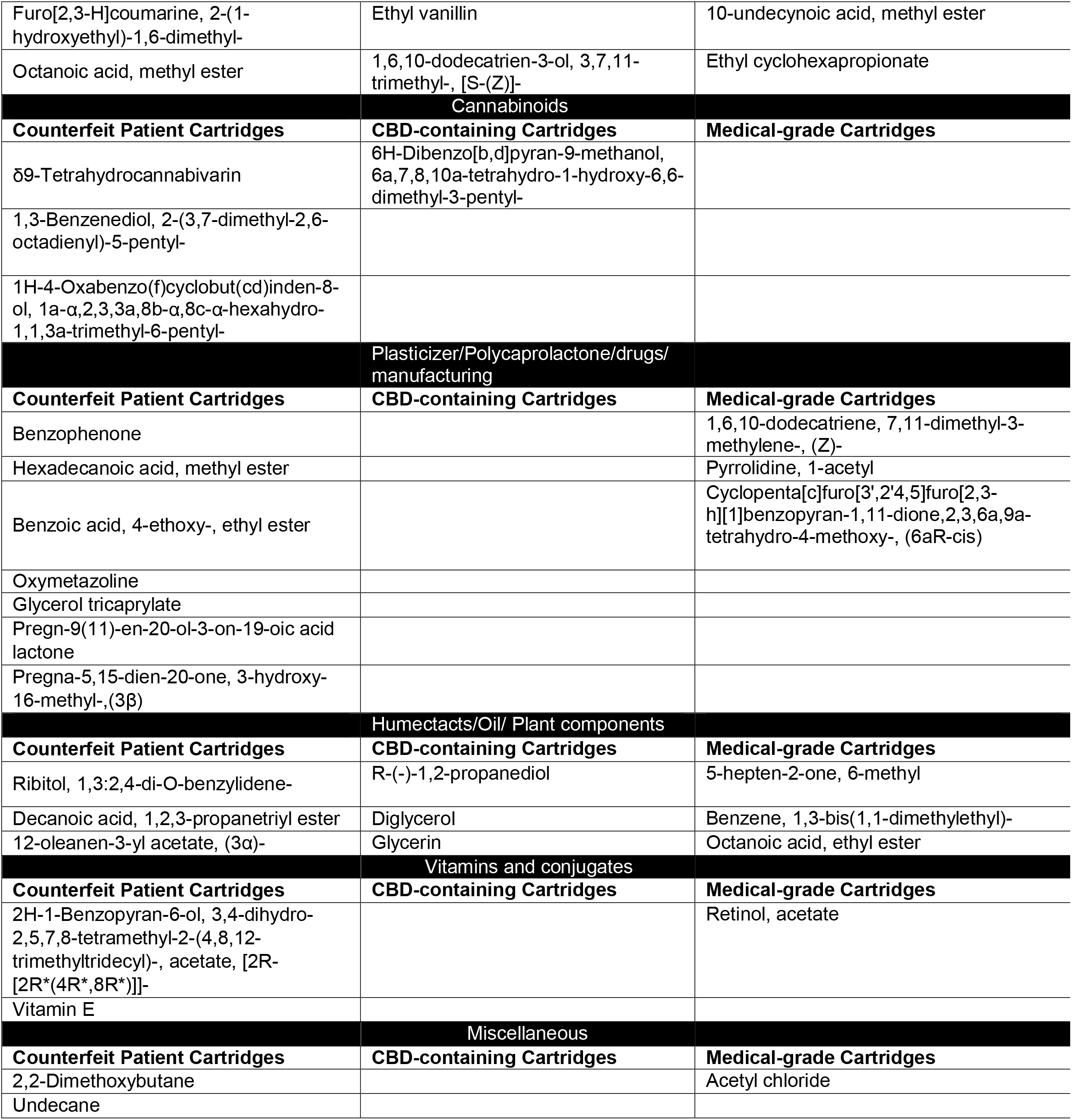
Commonly present constituents in counterfeit, CBD-containing, and medical-grade cartridges.

### Quantifying vapor phase chemical constituents

Aerosols from e-cig vaping cartridges were sampled in 1L vacuum bottles, and each cartridge was sampled for 10 minutes with 10 puffs each. These samples were sent to ALS Environmental, CA, for analysis. Vapor phase constituents, including VOC, were screened and quantified by EPA method TO-15 and mass spectral library search for tentatively identified compounds. Most abundant compounds were then classified into groups based on their use, i.e., terpenes, manufacturing/pesticides, automotive, solvents, PCP/household, and miscellaneous constituents **(Table 2)**.

**Table 2:**
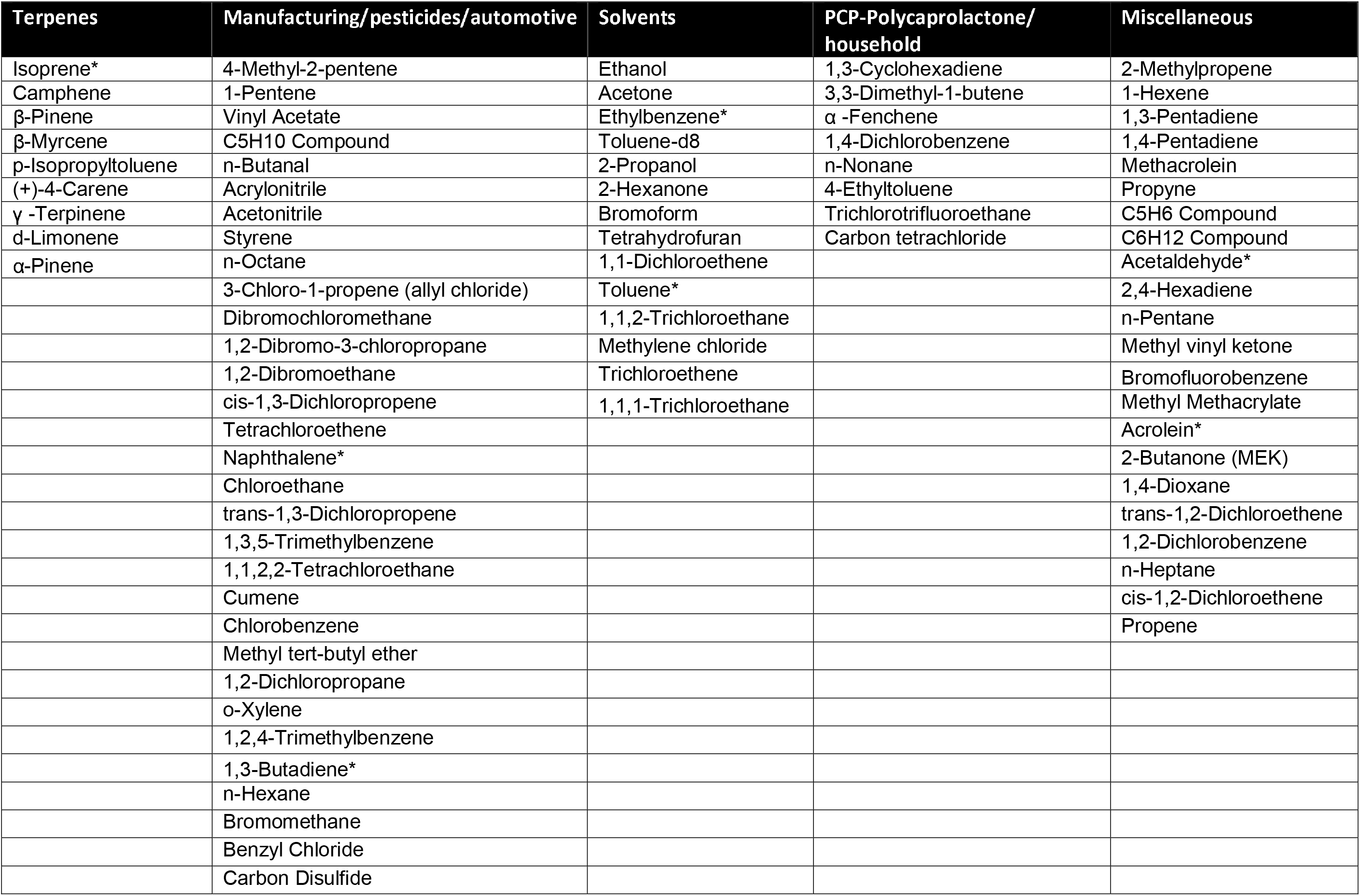
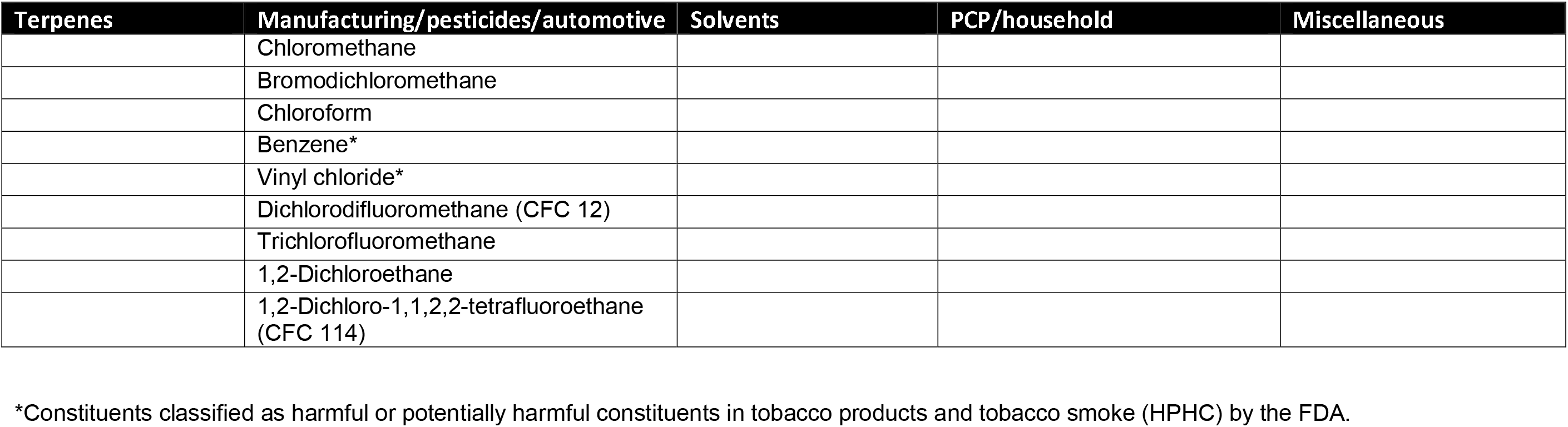
Vapor phase constituents in counterfeit cartridges.

### Elemental analysis of cartridge liquids

To screen for elements liquid aliquots from CBD-containing and counterfeit/patient-provided cartridges, ICP-MS analysis was performed. Total Quant KED analysis of elements used NeXion 2000 ICP-MS with external standards (20 ppb) solution in 2% nitric acid of 50 elements, adding helium gas at a rate of 4 mL/min. The solutions were pumped into a Meinhard nebulizer cooled to 2°C. This generated an aerosol that was aspirated into the plasma torch, where ionization occurs. The ion beam was then detected by the mass spectrometer. The most abundant elements in the liquids, particularly metals, were then tabulated **(Table 3)**.

**Table 3:**
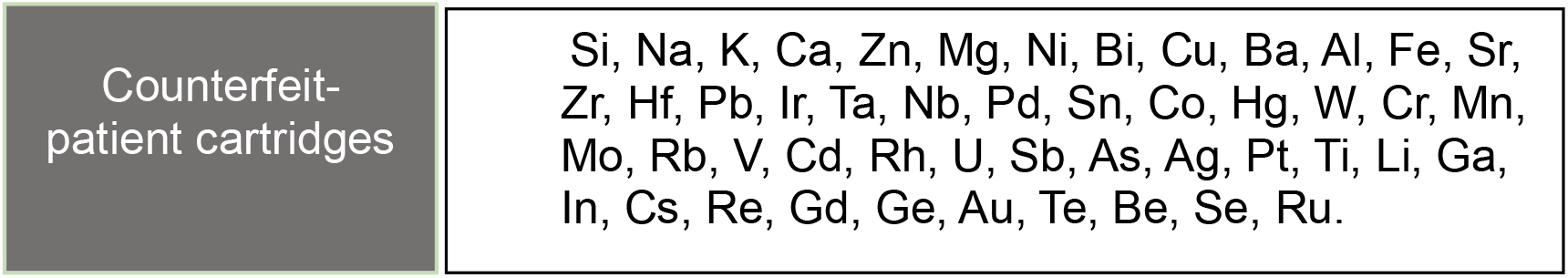
Elemental analysis of counterfeit cartridge liquids.

## Results

### Presence of unique chemicals in counterfeit patient-provided, CBD-containing, and medical-grade e-cig vaping cartridges

More than 500 chemicals were found in tested cartridges. Chemical constituents peculiarly present in e-cig counterfeit patient-provided, CBD-containing, and medical-grade cartridges are listed in **Table 1.** The majority of the compounds in e-cig cartridges were terpenes and silicon compounds. Chemicals unique to e-cig counterfeit patient-provided specific cartridges include 2,2-dimethoxybutane, tetramethyl silicate, decane, methyl esters, ethyl esters, siloxanes, and acetates, including α-tocopherol/vitamin E acetate (VEA). Apart from the chemicals listed above, e-cig counterfeit patient cartridges consisted of many terpenes and their acetates. Other compounds present included benzophenone, glycerol tricaprylate and similar products, and THC and its derivatives.

In most e-cig vaping CBD-containing cartridges, cannabidiol was present as opposed to THC. Glycerin, siloxanes, and flavoring chemicals, such as ethyl vanillin and coumarin compounds, were found in CBD-containing e-cig cartridges.

In e-cig medical-grade cartridges, acetyl chloride, vitamin A retinol-conjugated compounds, such as retinol acetate, rather than VEA, were present.

### Vapor-phase constituents in counterfeit e-cig cartridges

By screening for the vapor-phase constituents in counterfeit patient cartridges, we identified and quantified approximately 100 chemicals with concentrations ranging from 360,000 μg/m^3^ to 169 μg/m^3^ **(Table 2)**. Predominantly these vapor-phase constituents were manufacturing/pesticides/automotive/industrial active and inert agents and solvents.

### Metal constituents in e-liquids

Elemental analysis of the e-liquids from THC-containing counterfeit patient-provided cartridges was performed. Elements are presented in descending order, based on concentration found in the liquids **(Table 3)**. In counterfeit patient-cartridges, the most abundant elements included Si, Cu, Ni, and Pb. These metal concentrations were highly variable between cartridges, though the concentrations reached as high as 600 ppm.

## Discussion

In this study, we investigated the chemical constituents commonly found in e-cig vaping patient-provided counterfeit cartridges, CBD-cartridges, and medical cartridges. The presence of harmful compounds, such as MCT oil, VEA, and other lipids in THC-containing cartridges has been identified by the FDA/CDC (Blount et al. 2019a). At present, VEA has been linked as the causative agent based on bronchoalveolar lavage fluid (BALF) analyses in patients with EVALI (Blount et al. 2019b). Thus far, the EVALI epidemic has only been seen in North American populations, with very few cases reported in Canada (Stanbrook 2019). Interestingly, a recent study on e-cigarette ingredient analysis reported that there is no VEA in vaping products in the UK (Nyakutsikwa et al. 2019).

We determined the key constituents in counterfeit patient, CBD, and medical-grade e-cig vaping cartridges. We found several solvent chemicals, such as butane derivatives, i.e., 2,2 dimethoxybutane and n-butane, in both liquid and vapor phases of counterfeit e-cig cartridges. This is likely a result of “dabbing” with butane hash oil. Dabbing an extraction procedure commonly practiced in making illicit street cartridges as shown to improve total THC-recovery and lung availability by greater than 70%, which is unachievable by smoking cannabis alone (Hadener et al. 2019). Inhaling butane hash oil derivatives have already been seen as a probable cause for atypical/eosinophilic pneumonia in case reports (Anderson and Zechar 2019; McGraw et al. 2018).

The presence of hydrocarbons, such as decane and undecane, can also cause central nervous system damage, respiratory irritation, and even induce chemical pneumonitis (McKee et al. 2015). Similar to the effects of aspirating kerosene, these hydrocarbon solvents may induce pulmonary edema, cause lesions, destruct alveolar and capillary membranes, and alter surfactant production and function (Brown and Armstrong 2019). Indeed, the volatile organic compounds (VOC) were emitted higher in counterfeit patient cartridges (20.03±0.59) versus emitted by MCT (10.33±0.88), Mineral oil (7.33±0.88), and VEA (9.67±0.33, means ± SE) as monitored by QTrack VOC monitor. Further, their particle concentrations (mg/m^3^) in aerosols were also higher measured as particulate matter (diameter 1.0, 2.5, and 10 μm) in counterfeit patient cartridges vs MCT, mineral oil, and VEA, by Dustrack II 8530 instrument. Interactions between these hydrocarbons, particulates and other organic oils, such as mineral oil, MCT oil, and coconut oil, may promote or delay the absorption and metabolism of these compounds (Gerarde 1959). However, given the nature of the disease manifestation of the reported cases of EVALI with neutrophil and eosinophil infiltration in the BALF, and their successful treatment with steroids, most of the cases may have been chemical-induced pneumonitis (Henry et al. 2019; Triantafyllou et al. 2019).

In our chemical analysis, many conjugates and derivatives of silicon, such as silicates, silanes, and siloxanes, were identified only in e-cig counterfeit cartridges. Moreover, we also found large amounts of silica during elemental screening. These compounds, such as tetramethyl silicates, can form secondary products, such as silicon dioxide (SiO_2_) and methanol, which are highly toxic. Inhalation of silicon compounds is known to cause acute lung injury with pulmonary edema and lesions (NIOSH 2013).

It is believed that various hydrocarbons and reactive aldehydes are formed when e-liquids are heated to high temperatures around 500 °F. All these cartridges, including CBD containing cartridges, are used at the common voltages (e.g. 3.5V to 5.5 V) using a specific device (Chand et al. 2019). These conditions were used to generate aerosols for the detection of chemical constituents. Among the vapor-phase constituents, we found numerous known respiratory and cardiac toxicants specifically in e-cig counterfeit cartridges that are listed as harmful or potentially harmful constituents in tobacco products and tobacco smoke (HPHC) by the FDA. These compounds found in e-cig counterfeit cartridges include isoprene, acetaldehyde, ethylbenzene, toluene, acrolein, naphthalene, 1,3-butadiene, benzene, and vinyl chloride (FDA 2012).

Even though we detected VEA, we did not see VEA in all patient-provided counterfeit/street cartridges, only some cartridges contained VEA. Interestingly, medical-grade cartridges (formulation in coconut oil) contained retinol (vitamin A) acetate rather than VEA. Though there were still harmful chemicals, such as acetyl chloride, benzene, aflatoxin b1 identified in medical-grade cartridges, the number of contaminants detected in these cartridges were fewer compared to e-cig counterfeit patient-provided cartridges.

In counterfeit patient cartridges, we also found a common constituent in nasal sprays, oxymetazoline (decongestant), an α_1_ adrenergic receptor agonist. Common pesticide/insecticide ingredients, such as naphthalene and hexadecanoic, methyl ester were also detected in liquid and vapor phases of these e-cig counterfeit /patient cartridges. Exposure to very low levels of naphthalene, in both young and older mice, has been shown to induce acute cytotoxicity and injury to airway bronchiolar epithelial club cells (Carratt et al. 2019).

In the vapor-phase of e-cig counterfeit street cartridges, we detected many respiratory depressants and paralytic agents, such as 4-methyl-2 pentene, acetaldehyde, ethylbenzene, xylenes, and 1,3 pentadiene. Inhalation of these agents is likely the cause of symptoms, such as dyspnea and chest tightness (common symptoms among EVALI patients). Inhalation of compounds, such as acetone and trans -1,2, dichloroethane, and vinyl chloride, can cause drowsiness, which is another common symptom among EVALI patients. Among the vapor phase chemicals, inhalation of alkanes, such as n-butane and n-octane, causes acute lung toxicity. Moreover, exposure to toxicants such as methyl vinyl ketone and allyl chloride is highly toxic and causes emphysema, edema, and even vision impairment.

Chemical constituents in counterfeit cartridges, e.g., 1,3-butadiene, can self-polymerize and form peroxides upon exposure to oxygen. These chemicals may react with other compounds and further be catalyzed into forming secondary products (Levin et al. 2004). Inhaled xenobiotics undergo metabolism via phase I (CYP450/monooxygenase) and phase II (transferases) forming metabolites and detoxification/elimination products. During these processes, toxic metabolites may be formed. For example 1,3-butadiene has possible intermediates that are DNA reactive metabolites, such as butadiene monoepoxide and butadiene diepoxide (Dahl and Gerde 1994). Aside from the detected e-liquid chemical constituents, the lipid derivatives from the ‘endogenous’ source, such as the epithelial lining fluid (ELF) and/or lung surfactants by interacting with exogenous hydrocarbons with phospholipids and surfactants of the ELF, may also occur. This may be associated with the inflammatory responses using these counterfeit cartridges seen in the lungs of patients with EVALI (Chand et al. 2019). It remains to be seen whether the identified chemicals have any role for pathological changes in the lung, such as centrilobular ground-glass nodules and ground-glass opacities with subpleural sparing, as seen in patients with EVALI.

One of the limitations of this study includes the qualitative nature of the found constituents/toxicants in cartridge liquids as the GC-MS method established for this study was for preliminary screening to determine the presence of all constituents. It is possible that the other States may have different counterfeit cartridges involved in EVALI, which need to centralized for generating a library of chemicals identified in those cartridges for toxicological studies. Contributing susceptibility factors for EVALI pathogenesis may include genetic, environmental, lifestyle factors and concomitant diseases as not all THC-cartridge users were hospitalized for EVALI (Chatham-Stephens et al. 2019). The use of other vaping products, such as flavored e-liquids and pods with nicotine salts can be contributing and exacerbation factors to the pathogenesis of EVALI (Balmes 2020; Lu et al. 2020).

Legal restrictions on cannabis products limit research avenues to study the effects of these e-cig vaping products using surrogate models. Thus, there is a paucity of risk assessment and toxicological data on THC/cannabis containing products. Researchers and federal health organizations, such as the Centers for Disease Control (CDC) need to implement and adapt current e-cigarette research studies and clinical trials, and modify the established protocols to inform the participants and minimize the risk of developing EVALI.

In conclusion, our chemical analysis in e-cig counterfeit street/patient THC-containing cartridges showed numerous lethal respiratory toxicants in liquid and vapor phases, and the symptoms observed in patients are similar to what can be anticipated from inhaling those compounds. The potential toxicants include solvent derived hydrocarbons, silicon conjugated compounds, various terpenes, pesticides/plasticizers/polycaprolactones and metals. These chemicals are known to cause symptoms, such as cough, shortness of breath, or chest pain, nausea, vomiting, or diarrhea fatigue, fever, or weight loss as seen in patients with EVALI (Kalininskiy et al 2019). Our data suggest that exposure to a combination of hydrocarbons and oils, along with other toxic chemical compounds and metals, maybe a possible cause of EVALI as opposed to a singular causative agent. We are further investigating this using various *in vitro* and *in vivo* models before conclusively identifying the EVALI causative agent(s), as it may well be several agents involved in EVALI.

## Acknowledgments

Funding source: None

The authors thank Gary Ginsberg, Ph.D. at the New York State Department of Health, for his active involvement and input on our ongoing EVALI studies.

The authors thank Daniel Croft, MD, and Nicholas Nacca, MD at the Strong Memorial Hospital, Rochester for providing us information on hospitalized patient cartridges.

We thank Mr. Thomas Scrimale at the metal analysis core at the University of Rochester for performing the elemental analysis.

We thank Mr. Petar Nastoski, The College at Brockport, for preparing samples, performing GC/MS analysis and data processing.

We thank ALS Environmental, CA, for processing aerosol samples for vapor-phase constituents.

## Author contributions

Thivanka Muthumalage and Irfan Rahman conceived the study design, analyzed the data, wrote the manuscript, and edited the manuscript. Michelle Friedman and Alan Friedman performed the GCMS analysis and chemical identification, and edited the manuscript. Matthew McGraw and Michelle Friedman edited the manuscript.

